# CRISPR-Cas9 mediated mutagenesis of a *DMR6* ortholog in tomato confers broad-spectrum disease resistance

**DOI:** 10.1101/064824

**Authors:** Daniela Paula de Toledo Thomazella, Quinton Brail, Douglas Dahlbeck, Brian Staskawicz

**Affiliations:** Department of Plant and Microbial Biology, University of California, Berkeley, Berkeley CA 94720

**Keywords:** disease resistance, tomato, DMR6, CRISPR/Cas9 technology, crop engineering, food security

## Abstract

Pathogenic microbes are responsible for severe production losses in crops worldwide. The use of disease resistant crop varieties can be a sustainable approach to meet the food demand of the world’s growing population. However, classical plant breeding is usually laborious and time-consuming, thus hampering efficient improvement of many crops. With the advent of genome editing technologies, in particular the CRISPR-Cas9 (<u>c</u>lustered <u>r</u>egularly <u>i</u>nterspaced <u>s</u>hort <u>p</u>alindromic <u>r</u>epeats-Cas9) system, we are now able to introduce improved crop traits in a rapid and efficient manner. In this work, we genome edited durable disease resistance in tomato by modifying a specific gene associated with disease resistance. Recently, it was demonstrated that inactivation of a single gene called *DMR6* (*downy mildew resistance 6*) confers resistance to several pathogens in *Arabidopsis thaliana*. This gene is specifically up-regulated during pathogen infection, and mutations in the *dmr6* gene results in increased salicylic acid levels. The tomato *SlDMR6-1* orthologue Solyc03g080190 is also up-regulated during infection by *Pseudomonas syringae* pv. *tomato* and *Phytophthora capsici*. Using the CRISPR-Cas9 system, we generated tomato plants with small deletions in the *SlDMR6-1* gene that result in frameshift and premature truncation of the protein. Remarkably, these mutants do not have significant detrimental effects in terms of growth and development under greenhouse conditions and show disease resistance against different pathogens, including *P. syringae*, *P. capsici* and *Xanthomonas* spp.

## Introduction

Tomato (*Solanum lycopersicum*) is one of the most important crops and the second most consumed vegetable in the world. In particular, the United States is one of the world’s leading producers of tomatoes, accounting for a $2-billion-dollar annual market. Tomato is part of the Solanaceae family, which also includes many other economically important crops, such as pepper, potato and eggplant. Despite its great economic importance, plant pathogens (e.g. *Pseudomonas syringae*, *Phytophthora* spp., *Xanthomon*as spp.) are still a major limiting factor for tomato production around the world (Schwartz *et al.*, 2015). Plant breeding can be a sustainable and effective approach to obtain disease resistance in tomato. However, the classical plant-breeding methods are in general slow, laborious and time-consuming. Targeted genome editing has emerged as a promising alternative to classical breeding. This strategy employs programmable nucleases that enable a broad range of genetic modifications by inducing double-strand breaks in the DNA at specific genomic locations. These breaks stimulate error-prone non-homologous end joining (NHEJ) or homology-directed repair (HR) machineries. NHEJ repair can cause insertions or deletions (indels) around the DNA breaks, whereas HR can repair double-strand breaks using a homologous repair template (Joung and Sander, 2013; Liu and Fan, 2014; Urnov *et al.*, 2010).

Until recently, there were two types of programmable nucleases available for genome editing: the ZFNs (Zinc Finger Nucleases) and TALENs (Transcription Activator Like Effector Nucleases). These chimeric nucleases are composed of programmable, sequence-specific DNA-binding modules linked to a non-specific DNA cleavage domain (Joung and Sander, 2013; Urnov *et al.*, 2010). However, given the complexity and high costs for designing these proteins, these technologies have not been widely adopted by the plant research community. In the recent years, a novel precise tool for targeted genome editing was developed. It is based on the bacterial CRISPR-associated protein-9 nuclease (Cas9) from *Streptococcus pyogenes* (Liu and Fan, 2014). The CRISPR-Cas9 system is a bacterial immune system that has been modified for genome engineering. It is based on the Cas9 nuclease and an engineered single guide RNA (sgRNA) that specifies a targeted nucleic acid sequence. The target specificity relies on ribonucleotide-protein complex formation and not on protein-DNA interaction and thus gRNAs can be designed easily and economically to specifically target any sequence in the genome that is close to an NGG sequence [protospacer adjacent motif (PAM)], required for Cas9 recognition (Liu and Fan, 2014). Since the first application of the CRISPR-Cas9 system in mammalian cells (Cong *et al.*, 2013), it has been extensively used for genome editing in many organisms (Hwang *et al.*, 2013; Dicarlo *et al.*, 2013; Jiang *et al.*, 2013; Xie and Yang, 2013; Y., Wang *et al.*, 2013; Friedland *et al.*, 2013; Yu *et al.*, 2013; Nakayama *et al.*, 2013; Editor, 2014; D., Li *et al.*, 2013; Shen *et al.*, 2013; H., Wang *et al.*, 2013; W., Li *et al.*, 2013). In plants, the first uses of the CRISPR system were reported in 2013, with successful application for both transient expression and recovery of stable transgenic lines (J.,-F., Li *et al.*, 2013; Nekrasov *et al.*, 2013).

Over the last 25 years, the plant immune system has been extensively studied. These fundamental discoveries have led to a mechanistic understanding of the molecular basis of plant disease resistance and have enabled the identification of target host genes that can be altered by genome editing to produce disease resistant plants. In a recent report by Zeilmaker *et al.* (2015), it has been shown that mutation of a single gene associated with salicylic acid (SA) homeostasis results in the generation of Arabidopsis plants that are resistant to different types of pathogens, including bacteria and oomycetes. This gene is called *DMR6* (downy mildew resistance 6), and was originally identified and characterized by van Damme *et al*. (van Damme *et al.*, 2005; van Damme *et al.*, 2008). *DMR6* belongs to the superfamily of 2-oxoglutarate Fe(II) dependent oxygenases and is specifically up-regulated during pathogen infection. Interestingly, *dmr6* mutants have increased expression of defense genes and SA levels that correlate with enhanced resistance to *P. syringae*, *Hyaloperonospora arabidopsidis* and *Phytophthora capsici*. A *DMR6* paralog (*DLO1* or *DMR6*-like oxygenase), which is highly co-regulated with *DMR6*, was later identified in Arabidopsis and showed synergic effects with *DMR6* on plant resistance. *dlo1* mutants showed a lower level of resistance to *H. arabidopsidis* in relation to *dmr6* mutants, whereas *dmr6dlo1* double mutants showed complete resistance. Moreover, it was verified that increased SA levels are the basis of resistance, and these mutants require SA accumulation for the activation of plant immunity. Although the *dmr6dlo1* double mutant shows strong impaired growth presumably due to constitutive activation of immunity, the *dmr6* single mutant only shows slight growth reduction, which is possibly a consequence of a lower constitutive activation of plant defenses (Zeilmaker *et al.*, 2015). Considering this minimal effect on plant growth and the broad-spectrum resistance phenotype, generating *dmr6* knockouts is a promising strategy to control diseases in crop plants.

In this study, we identified a *DMR6* orthologue in the tomato genome and employed the CRISPR-Cas9 system to engineer disease resistance in this crop by inactivating this gene. We engineered tomato plants containing frameshift deletions in the *DMR6* gene that showed disease resistance against a wide variety of pathogens, but no significant effect on plant growth and development. Currently, the performance of these tomato varieties are being evaluated under conditions that are relevant to agricultural practices.

## Materials and methods

### Identification of tomato orthologues of *AtDMR6*

To search for *AtDMR6* homologous sequences in tomato, we performed a phylogenetic analysis using the following plant species: *Arabidopsis thaliana*, *Theobroma cacao, Manihot esculenta*, *Nicotiana benthamiana* and *Solanum lycopersicum*. Sequences were downloaded from the Phytozome database (https://phytozome.jgi.doe.gov/pz/portal.html) and Sol Genomics databases (https://solgenomics.net/). Proteins that contain the 2-oxoglutarate Fe(II) dependent oxygenase Pfam domain PF03171 were selected. The alignment was performed using Clustal Omega, and the tree was calculated using the neighbor-joining method. *AtDMR6* homologues were defined as those tomato genes that formed a clade along with *AtDMR6* (TAIR ID At5g24530).

### CRISPR-Cas9 construct design

**(A)** Specific gRNAs were designed to generate mutations within the coding sequence of the tomato *DMR6* gene Solyc03g080190 (exons 2 and 3). The gRNA sequences were assembled alongside the Cas9 endonuclease gene in the entry vector pENTR/D-TOPO. To drive the expression of both CRISPR components (gRNA and Cas9), we used the Arabidopsis U6-26 (AtU6-26) promoter and a 2 × 35S promoter, respectively, as previously described (Hwang *et al.*, 2013). gRNAs were assembled using the Golden Gate cloning method (Weber *et al.*, 2011). The insert containing the gRNA sequence and Cas9 gene is flanked by the attL1 and attL2 recombination sites and was gateway cloned into the binary destination vector pPZP200 as described by Katzen (Katzen, 2007).

### Transient assays in *Nicotiana benthamiana* for evaluation of gRNA efficiency

gRNA efficiency was first evaluated in transient expression assays using *Nicotiana benthamiana*. To generate the tomato target template in the living *N. benthamiana* cells, a fragment of about 1000 bp of *SlDMR6* was cloned into the binary geminivirus vector pLSLR to generate several copies of the target sequence. The gRNA/Cas9 pPZP200 and DMR6 template pLSL constructs were independently mobilized into *Agrobacterium tumefaciens* strain GV3101 by the triparental mating method (Wise *et al.*, 2006). The transformed *A. tumefaciens* cells (OD600 0.15) were syringe-infiltrated into *N. benthamiana* leaves for transient transformation. After four days, samples were collected for gDNA extraction, PCR analysis and amplicon sequencing with specific primers to detect mutations in the targeted regions of *SlDMR6* exons 2 and 3.

### *Agrobacterium*-mediated transformation of tomato

The binary vectors containing gRNAs 1 and 2 were mobilized into *A. tumefaciens* strain C58C1 by the triparental mating method (Wise *et al.*, 2006). *A. tumefaciens*-mediated transformations of the *S. lycopersicum* line FL8000 were performed according to Wang, 2015. In brief, cotyledon segments from 6-to 8-day-old seedlings were pre-cultured for one day followed by inoculation with *A. tumefaciens* strain C58C1 containing the CRISPR-Cas9 constructs of interest. Following a 2-day co-cultivation, the cotyledon segments were transferred to a selective regeneration medium containing 75 mg/l kanamycin. When shoots were approximately 1.5 cm tall, they were transferred to a selective rooting medium that also contained 75 mg/l kanamycin. Well-rooted plants were transferred to soil and grown for fruits production.

### Pathogen assays

FL8000 and *dmr6-1* tomato seeds (T1 generation) were planted in Sunshine Aggregate Mix #4 in 3 L plastic pots. Seedlings were grown in a glass house at temperatures ranging from 25 to 35°C. In planta assays were conducted in controlled environmental growth chambers to compare the growth curves of the *Sldmr6-1* mutants with the wild type parent strain FL8000. Bacterial cultures for plant inoculations were grown in NYGA media (peptone 5 g/L, yeast extract 3 g/L, agar 15 g/L, glycerol 20 ml/L) for 18 h at 28°C on a rotary shaker at 150 rpm. Cells were pelleted by centrifugation (4,000 × g, 15 min) and resuspended in sterile MgCl_2_ solution 10 mM. Bacterial suspensions were standardized to an optical density at 600 nm of 0.1 and subsequently used to infect the plants. Plants were inoculated with each species by being dipped into bacterial suspensions of *X. gardneri* (Xg153), *X. perforans* (Xp4b) and *P. syringae* pv. *tomato* (Pst DC3000) amended with the surfactant Silwet L-77 at 0.02% for 30 s. Following inoculation, plants were bagged and kept in a growth room on a 12-h photoperiod of fluorescent light at 24 to 28°C. Three samples were taken from each treatment after 6 (Pst), 12 (Xg) and 14 days (Xp). Bacterial populations were quantified by macerating 1-cm2 leaf disks in 1 ml sterile MgCl_2_ solution 10 mM, running 10-fold serial dilutions in sterile MgCl_2_ solution 10 mM, and plating them onto NYGA medium amended with rifampicin 100 μg/ml. After incubation at 28°C for 4 to 5 days, colonies typical of *Xanthomonas* spp./*P. syringae* were observed and counted. Data were log10 transformed, and standard errors were determined. Bacterial growth in the infected leaves of each plant was determined at days 6 post-inoculation (6 dpi), 12 post-inoculation (12 dpi) and 14 post-inoculation (14 dpi) for *P. syringae*, *X. gardneri* and *X. perforans*, respectively, according to the method of (Katagiri *et al.*, 2002).

For *P. capsici* pathogen assay, isolate LT1534 was grown on V8 agar 10% at 25°C for three days in the dark and for additional two days under fluorescent light. For inoculation, a plate covered with mycelium was flooded with cold water and the zoospore suspension was obtained after 30 minutes at room temperature. Leaves were spot-inoculated by pipetting 10 μl droplets of the spore suspension (10^5^ spores/ml) on the abaxial and adaxial sides.

### Measurement of tomato growth

To better determine the effects of DMR6 impairment on plant growth, the height of 28 *Sldmr6-1* mutants and 16 wild type plants was measured using a tape measure (Stanley FatMax 25’). The difference between the means was tested using the t-test at the significance level of p≤0.05.

## Results

### Introduction of CRISPR-Cas9 mutations into the *dmr6* gene of *Solanum lycopersicum*

DMR6 is part of the superfamily of 2-oxoglutarate Fe(II) dependent oxygenases. We performed a phylogeny analysis of the family of 2-oxoglutarate oxygenases (Pfam domain PF03171) including five different plant species: *Arabidopsis thaliana* (109 proteins), *Theobroma cacao* (109 proteins)*, Manihot esculenta* (136 proteins), *Nicotiana benthamiana* (110 proteins) and *Solanum lycopersicum* (142 proteins). We considered DMR6 homologues those genes that belong to the same clade of *AtDMR6*, and two copies of the *DMR6* gene were identified in tomato (SlDMR6-1: Solyc03g080190 and SlDMR6-2: Solyc06g073080) (Figure 1).

**Figure 1:**
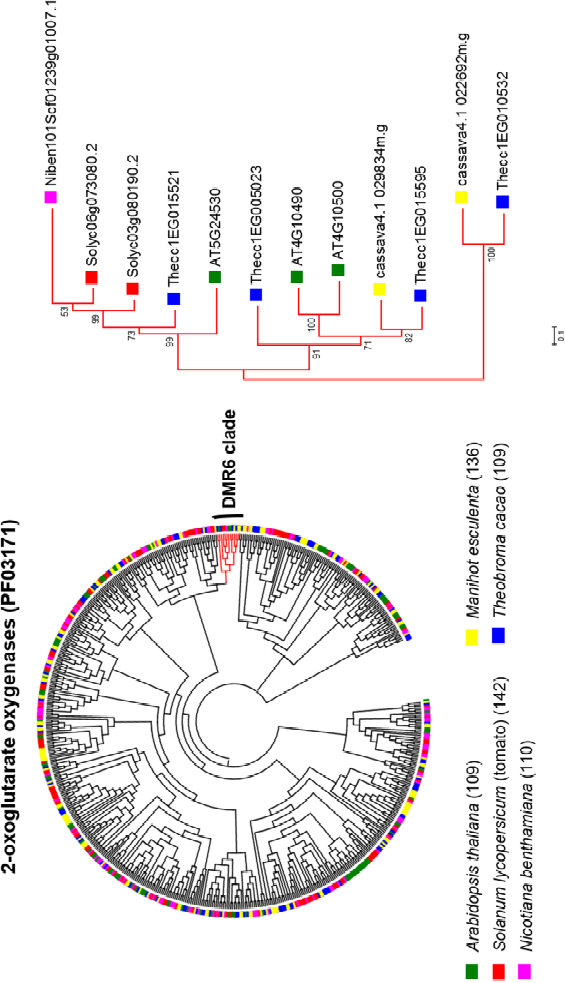
Phylogenetic tree of 2-oxoglutarate oxygenase proteins from *Arabidopsis thaliana* and four other plants (left panel), including the Solanaceae model *Nicotiana benthamiana* and three crops (*Solanum lycopersicum, Manihot esculenta* and *Theobroma cacao*). The DMR6-clade of 2OG oxygenases is highlighted on the right. Bootstrap values are shown in the tree.

Based on the inspection of public transcriptomic data (Yang *et al.*, 2015; Jupe *et al.*, 2013), we verified that Solyc03g080190 is up-regulated by infection with *P. syringae* pv. *tomato* (DC3000) and *P. capsici*, which suggests that Solyc03g080190 might have a similar function to the Arabidopsis *DMR6* (Figure 2).

**Figure 2:**
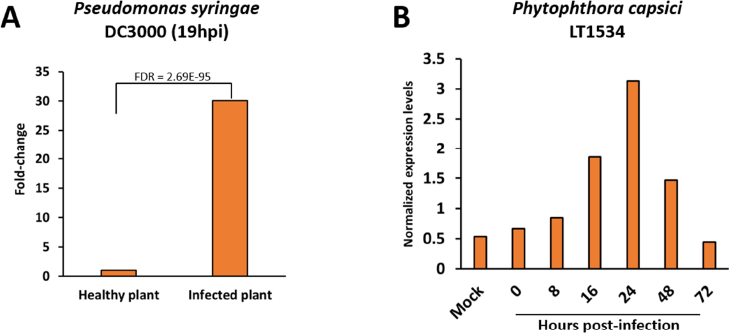
Expression of the tomato *DMR6* gene obtained from public transcriptomic data (Jupe *et al.*, 2013, Yang *et al.*, 2015) in response to pathogen infection. **A)** Tomato *DMR6* is upregulated at 19hpi by the bacteria *Pseudomonas syringae* pv. *tomato* (DC3000). NI= non-inoculated; hpi= hours after infection. **B)** Tomato *DMR6* is also upregulated by infection with the oomycete *Phytophthora capsici*.

We designed specific gRNAs that target exons 2 (target region: 5’-TAGAGAAGTATGCTCCTGAA-3’) and 3 (target region: 5’-AGTTCTGGTTGTGGACAAGG-3’) of the tomato *DMR6* gene Solyc03g080190. As gRNA efficiency can vary according to the gRNA sequence, before proceeding to tomato transformation, we tested gRNA efficiency using transient expression assays of the CRISPR constructs in *Nicotiana benthamiana*. We performed PCR analysis and amplicon sequencing with specific primers (5’-ATGGTGTACCAAAGGAAGTTGTAGAGA-3’ and 5’-TGCAACACTTCTCAGTTTGAGCCTCG-3’ for exon 2; 5’-AGATATTGCAGGGAAATTCGTCAACTC-3’; 5’-GATGCCATACACTTCTGTACTTACCGTT-3’ for exon 3) and detected mutations in the targeted regions of both gRNAs (Figure 3).

**Figure 3:**
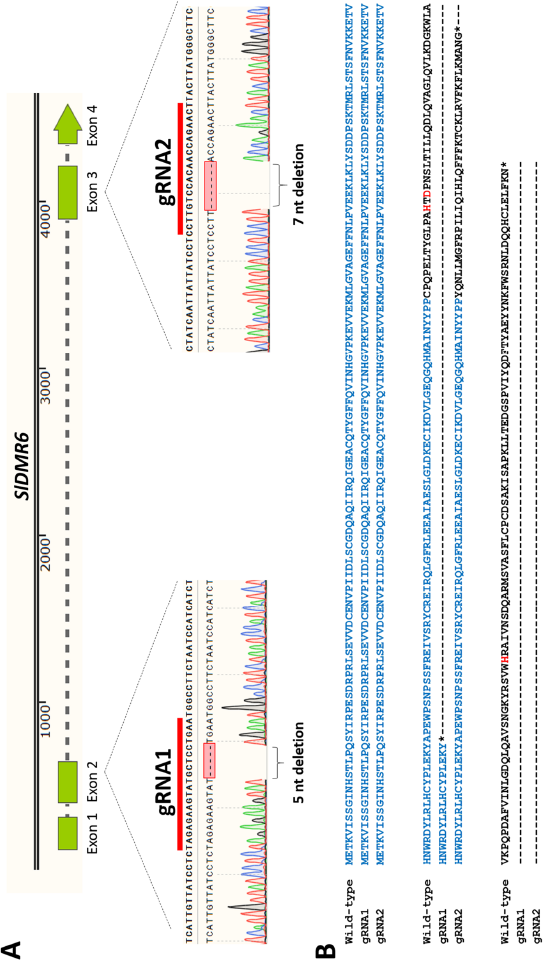
Specific gRNAs were designed to generate mutations within the coding sequence of the tomato *DMR6* gene Solyc03g080190.2.1 (exons 2 and 3). The gRNA target sequence is represented by a red bar over the gene sequence. Frameshift mutations (5 bp deletion in exon 2 and 7 bp deletion in exon 3) generated by these gRNAs are shown in details. These sequences are from homozygous mutants obtained in the T0 tomato plants. **(B)** Clustal aligment of the SLDMR6-1 amino acid sequences that correspond to the wild type line, the 5bp-deletion mutant (generated with gRNA1) and the 7bp-deletion mutant (generated with gRNA2) lines. The region highlighted in blue are the amino acids present in all three lines, and the amino acid residues highlighted in red are the conserved amino acids HxD/H that form the protein active site. These amino acids are essential for the protein enzymatic activity and are not present in the mutant lines described above.

Once we confirmed gRNA efficiencies, tomato plants (FL8000 line) were transformed via Agrobacterium with the binary vector containing one of the two gRNAs and the *Cas9* gene. After about 6 weeks, whole plants were successfully regenerated.

### Molecular characterization of *SlDMR6* mutations and transmission to F1 progeny

The CRISPR transformants were genotyped (PCR and sequencing of the target region) using the primers described above for the *N. benthamiana* transient assays. Different mutations in the *SlDMR6-1* gene were obtained, including frameshift deletions and insertions that truncate the protein and disrupt the DMR6 active site. As previously discussed, DMR6 belongs to the superfamily of 2-oxoglutarate Fe(II) dependent oxygenases. The active site of these enzymes is characterized by the conserved motif HxD/Ex H (Clifton *et al.*, 2006), and these amino acid residues are required for DMR6-dependent susceptibility phenotype in *A. thaliana* (Zeilmaker *et al.*, 2015). Remarkably, among the tomato mutant lines obtained, homozygous *Sldmr6-1* mutants were found in the primary transformants. Given that primary transformants can show somaclonal variation, seeds of the tomato *dmr6* mutants were obtained and the T1 plants were genotyped to be used in infection assays. In addition, we identified segregating progeny in the T1 tomato plants that maintain mutations in the *DMR6* gene and no longer contain the T-DNA with the Cas9 gene. (Figure 4).

**Figure 4:**
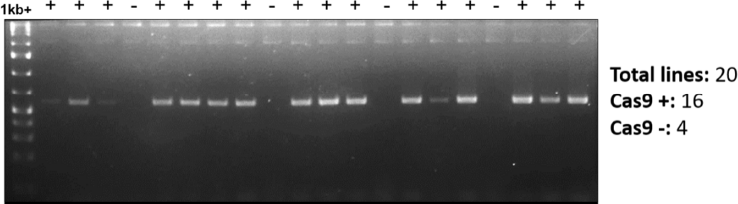
PCR with *Cas9* gene specific primers using gDNA extracted from *dmr6* tomato mutants (T1). In this case, the *Cas9* gene probably integrated in a single locus in the tomato genome and, as a consequence, the gene segregated out in about 20% of the T1 plants. The mutation in the *DMR6* gene is a 7 nucleotide deletion in exon 3 that disrupts the protein active site. + Cas9 is present; - Cas9 is absent.

### Characterization of growth traits and disease phenotypes of *Sldmr6-1* mutants

T1 plants derived from a homozygous line with a 7 bp deletion in exon 3 of the *DMR6* gene were used in infection assays with *Xanthomonas gardneri* (Xg153), *Xanthomonas perforans* (Xp4b), and *P. syringae* pv. *tomato* (DC3000). The bacterial growth assays were performed using the dip inoculation method. Overall, the tomato *dmr6* mutants were more resistant to all these pathogens, and the disease symptoms were less severe in comparison to the wild type lines (Figure 5). In addition, the oomycete pathogen *P. capsici* has also been tested, and it was able to infect both wild type and *dmr6* lines, but symptoms developed earlier and were more severe in the wild type plants (Figure 6).

**Figure 5:**
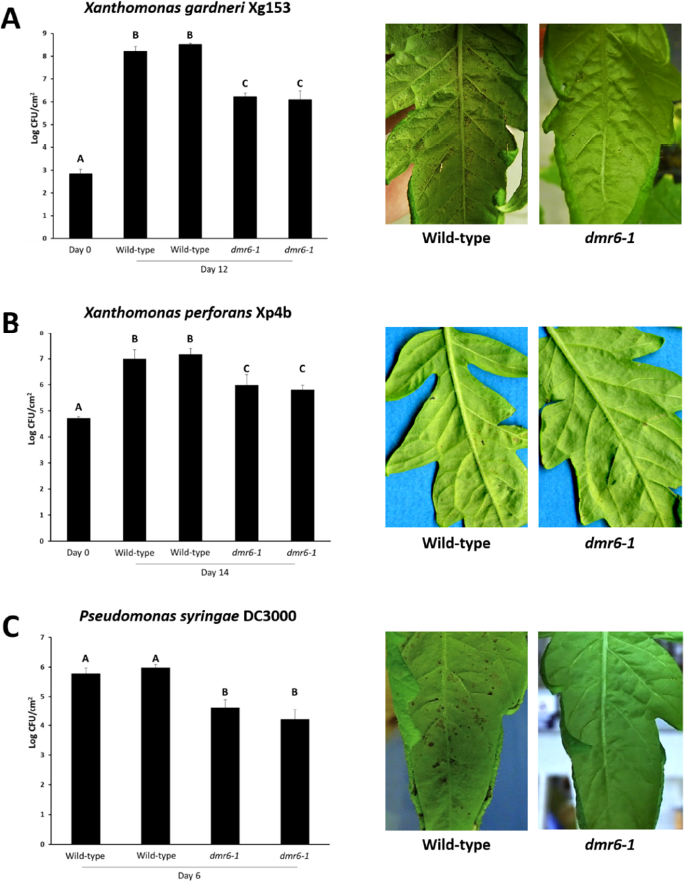
Wild type and *dmr6* mutant lines infected with *Xanthomonas gardneri* (Xg153)*, Xanthomonas perforans* (Xp4B) and *Pseudomonas syringae* pv. *tomato* (DC3000). On the left, bacterial counts for *X. gardneri* **(A)**, *X. perforans* **(B)** and *P. syringae* **(C)** are shown in the wild type and in the *dmr6* mutant lines at 0 dai (days after infection) and 12 dai, 14 dai and 6 dai, respectively. Inactivation of DMR6 results in lower bacterial numbers for all three species. Bars represent standard error of three replicates. On the right, lesions (black dots) in the leaf correspond to the disease symptoms, which are more severe in the wild type plants.

**Figure 6:**
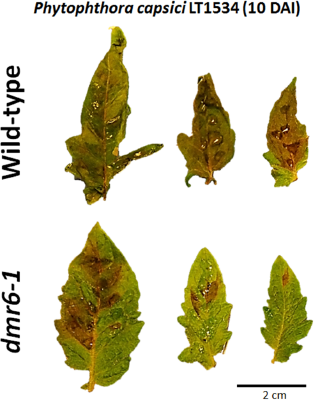
Wild type and *dmr6* mutant lines infected with *Phytophthora capsici* at 10 days after infection (dai). The necrosis in the leaf corresponds to the disease symptom, which is considerably more severe in the wild type plant. Inoculation was performed using droplets of a suspension of *P. capsici* zoospores at 10^5^ zoospores/ml.

To better determine the effects of DMR6 impairment on plant growth, the height of 28 *Sldmr6-1* mutants and 16 wild type plants was measured, and only a slight decrease in size, which was not statistically significant, was verified for the *Sldmr6-1* mutants (Figure 7).

**Figure 7:**
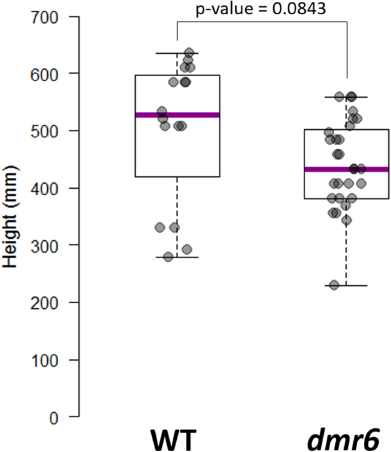
Height growth of tomato plants (*Sldmr6-1* and wild type lines). Height was measured along the length of the stem starting on the stem aboveground to the apex. Twenty-eight *Sldmr6-1* mutants and 18 wild type plants were used. The difference between the means was tested using the t-test at the significance level of p≤0.05.

## Discussion

One of the current challenges for food security is to increase crop yields and support a sustainable agriculture. Notably, pathogen infection is a major cause of crop yield losses around the world. Introgression of resistance (R) genes is the classical approach to achieve disease resistance in crops. This type of resistance leads to the development of the hypersensitive reaction (HR), which is a localized cell death triggered by the interaction between a plant R gene and its respective pathogen *avr* (avirulence) gene (Jones and Dangl, 2006). In addition to the use of plant R genes for resistance breeding, impairment of some plant genes via loss-of-function mutations has been shown to confer broad-spectrum disease resistance, as it is the case of the *A. thaliana DMR6* gene. In this study, we show that inactivation of the *AtDMR6* homologue in tomato results in resistance to different pathogens, such as the bacterial pathogens *P. syringae* and *Xanthomonas* spp. and the oomycete pathogen *P. capsici*.

DMR6 is part of the 2-oxoglutarate Fe(II)-dependent oxygenases superfamily, which is present in all angiosperms. Members of this superfamily are involved in secondary metabolism and biosynthesis of plant hormones (Zeilmaker *et al.*, 2015). The phylogenetic analysis of 2-oxoglutarate oxygenases showed the presence of two *AtDMR6* homologues in tomato, which grouped in the same clade of *AtDMR6* and were named *SlDMR6-1* and *SlDMR6-2* (Figure 1). These DMR6 homologues were analyzed in more detail since they could play a similar role in defense as *AtDMR6*. To better characterize their participation in plant defense, we first analyzed public transcriptomic data of tomato infected by *P. syringae* pv. *tomato* (DC3000) and *P. capsici*. Only *SlDMR6-1* was upregulated by pathogen attack (Figure 2), which suggests that this paralogue is important for plant immunity. Therefore, we focused on mutating the *SlDMR6-1* gene in tomato and verifying if impairment of this gene leads to broad spectrum disease resistance as it has been described for *A. thaliana*.

Remarkably, tomato *Sldmr6-1* mutants showed enhanced disease resistance to important bacterial pathogens of tomato, such as *P. syringae, X. gardneri and X. perforans* as well as to the oomycete pathogen *P. capsici* (Figures 5 and 6). These results indicate that the biological function of *AtDMR6* homologues might be conserved among different plant species. In particular, resistance to *Xanthomonas spp.* is highly desired. *X. gardneri and X. perforans* belong to a complex of four *Xanthomonas* species (*X. euvesicatoria, X. vesicatoria, X. perforans* and *X. gardneri*) that incite bacterial spot disease in tomato, one of the most devastating and widespread diseases of this crop that can cause significant losses when environmental conditions are suitable for the pathogen (i.e. moderate-to-high temperatures and high humidity) (Schwartz *et al.*, 2015). The use of chemicals for controlling bacterial spot disease have not been effective due to the emergence of tolerant strains and the significant negative impact on the environment (Abbasi *et al.*, 2015). For this reason, resistance against bacterial spot has been a priority in tomato breeding programs. The R protein *Bs2* has been identified in pepper, and is able to recognize a highly conserved Xanthomonas effector named *AvrBs2*, thus triggering HR. Field trials have shown that transgenic tomato plants expressing the *Bs2* gene show high disease resistance and significant increased yield (Horvath *et al.*, 2012). Although very promising, resistance based on the deployment of a single R gene can be rapidly overcome by the pathogen due to the emergence of resistant strains (Dangl *et al.*, 2013). Moreover, *Bs2* specifically recognizes a single effector in a specific species, limiting the use of these resistant tomato lines. In this regard, to engineer long-lasting and broad-spectrum disease resistance in tomato, the combination of Bs2 deployment with DMR6 inactivation is a very promising approach.

Preliminary data indicate that SA levels are increased in the *Sldmr6-1* mutants (data not shown) in comparison to the wild type tomato plants, suggesting that, resistance correlates with increased SA levels. SA is a central phytohormone in defense responses against biotrophic and hemibiotrophic pathogens. It induces plant production of secondary metabolites as well as expression of pathogenesis-related protein genes. Also, SA-mediated responses often culminate with the onset of HR. Interestingly, mutants displaying a broad-spectrum disease resistance phenotype derived from increased SA levels have already been described. The *cpr* (*constitutive expressor of PR genes*) mutants exhibit increased concentrations of SA, constitutive expression of PR genes, and enhanced resistance to different pathogens. However, these mutants have a severe dwarf phenotype, which is a consequence of the high-energy expenditure on keeping the defense responses active. Importantly, the tomato *Sldmr6-1* mutants did not show a significant detrimental effect on plant growth and development (Figure 7). This characteristic definitely makes DMR6 a very promising target for resistance breeding in crops. Future work will also evaluate the performance of these tomato *Sldmr6-1* mutants in field conditions.

Although it is still unclear how impairment of DMR6 results in increased salicylic acid levels, *AtDMR6* activity has been recently demonstrated by Falcone Ferreyra *et al.* (2015). The authors showed that *AtDMR6* as well as its ortologue in maize have flavone synthase activity and catalyze the conversion of the flavanone naringenin into the flavone apigenin (Falcone Ferreyra *et al.*, 2015). Flavonoids and SA pathways share some precursors, thus the authors suggest that when DMR6 is inactivated, a minor flow through the flavonoid pathway would lead to higher availability of substrates for SA biosynthesis, thus increasing the levels of SA. Moreover, the authors speculate that this would be a strategy used by pathogens to decrease SA levels and hence increase susceptibility (Falcone Ferreyra *et al.*, 2015). However, this hypothesis still needs to be confirmed.

Given the broad-spectrum disease resistance phenotype and the lack of significant detrimental effects in the plant, impairment of some plant genes to obtain disease resistance has been considered a promising strategy by many scientists in the field of plant pathology. Nevertheless, as the fitness costs of DMR6 inactivation have not been clearly determined yet, more studies are still needed to evaluate the performance of these mutant lines under conditions that are important to agricultural practices.

## Acknowledgements

We thank the 2Blades Foundation for the financial support and the Tom Clemente Lab (Nebraska University) for tomato transformation. DPTT is funded by the PEW Fellowship Program in the Biomedical Sciences. We also thank Paulo José Teixeira for careful reading of the manuscript.

## Author contribution

D.P.T.T. designed and performed experiments, analyzed data and wrote the manuscript; Q.B. and D.D. performed experiments; and B.J.S. supervised the project, designed the study and the experiments and analyzed data.

## References

Abbasi, P.A., Khabbaz, S.E., Weselowski, B. and Zhang, L. (2015) Occurrence of copper-resistant strains and a shift in Xanthomonas spp. causing tomato bacterial spot in Ontario. Can. J. Microbiol., 761, 1–9.

Clifton, I.J., McDonough, M.A., Ehrismann, D., Kershaw, N.J., Granatino, N. and Schofield, C.J. (2006) Structural studies on 2-oxoglutarate oxygenases and related double-stranded ??-helix fold proteins. J. Inorg. Biochem., 100, 644–669.

Cong, L., Ran, F.A., Cox, D., Lin, S., Barretto, R., Hsu, P.D., Wu, X., Jiang, W. and Marraffini, L.A. (2013) Multiplex Genome Engineering Using CRISPR/VCas Systems. Science (80-.)., 339, 819–823.

Damme, M. van, Andel, A., Huibers, R.P., Panstruga, R., Weisbeek, P.J. and Ackerveken, G. Van den (2005) Identification of arabidopsis loci required for susceptibility to the downy mildew pathogen Hyaloperonospora parasitica. Mol. Plant. Microbe. Interact., 18, 583–592.

Damme, M. van, Huibers, R.P., Elberse, J. and Ackerveken, G. Van Den (2008) Arabidopsis DMR6 encodes a putative 2OG-Fe(II) oxygenase that is defense-associated but required for susceptibility to downy mildew. Plant J., 54, 785−793.

Dangl, J.L., Horvath, D.M. and Staskawicz, B.J. (2013) Pivoting the plant immune system from dissection to deployment. Science, 341, 746–51. Available at: http://www.pubmedcentral.nih.gov/articlerender.fcgi?artid=3869199&tool=pmcentrez&rendertype=abstract.

Dicarlo, J.E., Norville, J.E., Mali, P., Rios, X., Aach, J. and Church, G.M. (2013) Genome engineering in Saccharomyces cerevisiae using CRISPR-Cas systems. Nucleic Acids Res., 41, 4336–4343.

Editor, D. (2014) Letter to the Editor Effective gene targeting in rabbits using RNA-guided Cas 9 nucleases., 97–99.

Friedland, A.E., Tzur, Y.B., Esvelt, K.M., Colaiacovo, M.P., Church, G.M. and Calarco, J.A. (2013) Heritable genome editing in C. elegans via a CRISPR-Cas9 system. Nat Methods, 10, 741–743.

Horvath, D.M., Stall, R.E., Jones, J.B., Pauly, M.H., Vallad, G.E., Dahlbeck, D., Staskawicz, B.J. and Scott, J.W. (2012) Transgenic resistance confers effective field level control of bacterial spot disease in tomato. PLoS One, 7.

Hwang, W.Y., Fu, Y., Reyon, D., Maeder, M.L., Tsai, S.Q., Sander, J.D., Peterson, R.T., Yeh, J.R. and Joung, J.K. (2013) Efficient genome editing in zebrafish using a CRISPR-Cas system. Nat Biotechnol, 31, 227–229. Available at: http://www.ncbi.nlm.nih.gov/pubmed/23360964\nhttp://www.nature.com/nbt/journal/v31/n3/pdf/nbt.2501.pdf.

Jiang, W., Zhou, H., Bi, H., Fromm, M., Yang, B. and Weeks, D.P. (2013) Demonstration of CRISPR/Cas9/sgRNA-mediated targeted gene modification in Arabidopsis, tobacco, sorghum and rice. Nucleic Acids Res., 41, 1–12.

Jones, J.D.G. and Dangl, J.L. (2006) The plant immune system. Nature, 444, 323–329.

Joung, J.K. and Sander, J.D. (2013) TALENs: a widely applicable technology for targeted genome editing. Nat Rev Mol Cell Biol, 14, 49–55. Available at: http://www.ncbi.nlm.nih.gov/pubmed/23169466\nhttp://www.nature.com/nrm/journal/v14/n1/pdf/nrm3486.pdf.

Jupe, J., Stam, R., Howden, A.J.M., Morris, J. a, Zhang, R., Hedley, P.E. and Huitema, E. (2013) Phytophthora capsici-tomato interaction features dramatic shifts in gene expression associated with a hemi-biotrophic lifestyle. Genome Biol., 14, R63.

Katagiri, F., Thilmony, R. and He, S.Y. (2002) The Arabidopsis Thaliana-Pseudomonas Syringae Interaction. Arab. B., 1, e0039. Available at: http://www.bioone.org/doi/abs/10.1199/tab.0039.

Katzen, F. (2007) Recombinational Cloning: a Biological Operating System. Expert Opin. Drug Discov., 2, 571–589.

Li, D., Qiu, Z., Shao, Y., et al. (2013) Heritable gene targeting in the mouse and rat using a CRISPR-Cas system. Nat. Biotechnol., 31, 681–683.

Li, J.-F., Norville, J.E., Aach, J., McCormack, M., Zhang, D., Bush, J., Church, G.M. and Sheen, J. (2013) Multiplex and homologous recombination–mediated genome editing in Arabidopsis and Nicotiana benthamiana using guide RNA and Cas9. Nat. Biotechnol., 31, 688–691. Available at: http://dx.doi.org/10.1038/nbt.2650\npapers2://publication/doi/10.1038/nbt.2650\nhttp://www.nature.com/doifinder/10.1038/nbt.2654.

Li, W., Teng, F., Li, T. and Zhou, Q. (2013) Simultaneous generation and germline transmission of multiple gene mutations in rat using CRISPR-Cas systems. Nat. Biotechnol., 31, 684–686.

Liu, L. and Fan, X.-D. (2014) CRISPR--Cas system: a powerful tool for genome engineering. Plant Mol. Biol., 85, 209–218. Available at: http://dx.doi.org/10.1007/s11103-014-0188-7.

Nakayama, T., Fish, M.B., Fisher, M., Oomen-Hajagos, J., Thomsen, G.H. and Grainger, R.M. (2013) Simple and efficient CRISPR-Cas9-mediated targeted mutagenesis in Xenopus tropicalis. Genesis, 51, 835–843.

Nekrasov, V., Staskawicz, B., Weigel, D., Jones, J.D.G. and Kamoun, S. (2013) Targeted mutagenesis in the model plant Nicotiana benthamiana using Cas9 RNA-guided endonuclease. Nat Biotech, 31, 691–693. Available at: http://dx.doi.org/10.1038/nbt.2655.

Schwartz, A.R., Potnis, N., Timilsina, S., et al. (2015) Phylogenomics of Xanthomonas field strains infecting pepper and tomato reveals diversity in effector repertoires and identifies determinants of host specificity. Front. Microbiol., 6. Available at: http://journal.frontiersin.org/Article/10.3389/fmicb.2015.00535/abstract [Accessed July 12, 2016].

Shen, B., Zhang, J., Wu, H., Wang, J., Ma, K., Li, Z., Zhang, X., Zhang, P. and Huang, X. (2013) Generation of gene-modified mice via Cas9/RNA-mediated gene targeting. Cell Res., 23, 720–3.

Urnov, F.D., Rebar, E.J., Holmes, M.C., Zhang, H.S. and Gregory, P.D. (2010) Genome editing with engineered zinc finger nucleases. Nat Rev Genet, 11, 636–646. Available at: http://www.ncbi.nlm.nih.gov/pubmed/20717154\nhttp://www.nature.com/nrg/journal/v11/n9/pdf/nrg2842.pdf.

Wang, H., Yang, H., Shivalila, C.S., Dawlaty, M.M., Cheng, A.W., Zhang, F. and Jaenisch, R. (2013) One-step generation of mice carrying mutations in multiple genes by CRISPR/cas-mediated genome engineering. Cell, 153, 910–918.

Wang, K. (2015) Methods in Molecular Biology, vol 343: Agrobacterium protocols-vol I, Available at: www.humanapress.com.

Wang, Y., Li, Z., Xu, J., Zeng, B., Ling, L., You, L., Chen, Y., Huang, Y. and Tan, A. (2013) The CRISPR/Cas system mediates efficient genome engineering in Bombyx mori. Cell Res., 23, 1414–1416.

Weber, E., Engler, C., Gruetzner, R., Werner, S. and Marillonnet, S. (2011) A modular cloning system for standardized assembly of multigene constructs. PLoS One, 6.

Wise, A. a, Liu, Z. and Binns, A.N. (2006) Three methods for the introduction of foreign DNA into Agrobacterium. Methods Mol. Biol., 343, 43–53. Available at: http://www.ncbi.nlm.nih.gov/pubmed/16988332.

Xie, K. and Yang, Y. (2013) RNA-Guided genome editing in plants using a CRISPR-Cas system. Mol. Plant, 6, 1975–1983.

Yang, Y.-X., Wang, M.-M., Yin, Y.-L., et al. (2015) RNA-seq analysis reveals the role of red light in resistance against Pseudomonas syringae pv. tomato DC3000 in tomato plants. BMC Genomics, 16, 120. Available at: http://www.pubmedcentral.nih.gov/articlerender.fcgi?artid=4349473&tool=pmcentrez&rendertype=abstract.

Yu, Z., Ren, M., Wang, Z., Zhang, B., Rong, Y.S., Jiao, R. and Gao, G. (2013) Highly efficient genome modifications mediated by CRISPR/Cas9 in Drosophila. Genetics, 195, 289–291.

Zeilmaker, T., Ludwig, N.R., Elberse, J., Seidl, M.F., Berke, L., Doorn, A. Van, Schuurink, R.C., Snel, B. and Ackerveken, G. Van Den (2015) DOWNY MILDEW RESISTANT 6 and DMR6-LIKE OXYGENASE 1 are partially redundant but distinct suppressors of immunity in Arabidopsis., 210–222.

